# An Optimized Stem Cell Secretome Proteomics Platform: Application to Progranulin-Deficient iPSCs

**DOI:** 10.64898/2026.08.18.745538

**Authors:** Jiawei Ni, Hope Tracey, Ling Hao

**Author notes:** Corresponding author Ling Hao, PhD. Associate Professor, Department of Chemistry & Biochemistry University of Maryland.

## Abstract

Stem cells secrete diverse extracellular proteins that regulate pluripotency, differentiation, and cell–cell communication, making them powerful model systems for studying development, disease mechanisms, and regenerative medicine. However, robust stem cell secretome analysis remains technically challenging. Unlike many other cell types, stem cells cannot tolerate serum starvation or growth factor deprivation, while low-abundance secreted proteins are often masked by media-derived proteins and intracellular contamination. Here, we systematically optimized the secretome proteomics workflow in iPSCs, by evaluating culture medium composition, conditioned-media collection time, cell plating density, media harvest and preparation methods, LC–MS acquisition methods, and data analysis strategies. Full-strength Essential 8 medium, 48 h media collection, 80% cell confluency, two-step centrifugation, and data-independent acquisition (DIA)-LC-MS/MS provided the optimal secretome proteomics data quality. We then applied the optimized platform to an isogenic iPSC disease model to investigate how progranulin deficiency reshapes the extracellular and intracellular proteomes. Progranulin-deficient iPSCs showed a coordinated reduction of extracellular lysosomal hydrolases despite relatively modest intracellular proteome changes, suggesting altered lysosome trafficking and possible impairment of lysosomal exocytosis. Together, this work establishes a robust and standardized workflow for stem cell secretome proteomics and demonstrates its utility for investigating extracellular proteome remodeling in human disease models.

## 1. Introduction

Stem cells secrete a broad spectrum of molecules, including growth factors, cytokines, extracellular matrix (ECM) components, exosome proteins, and enzymes, collectively shaping the extracellular signaling environment to regulate pluripotency, differentiation, and tissue repair^1–3^. In particular, human induced pluripotent stem cells (iPSCs) have become indispensable models for studying development, disease mechanisms, and regenerative medicine^4^. During early development, the iPSC secretome plays a central role in regulating cell–cell communication and remodeling the extracellular microenvironment to direct lineage commitment^5–7^. Because secreted proteins reflect ongoing signaling and extracellular remodeling events, secretome profiling provides a functional snapshot of how cells interact with and respond to their surroundings. Therefore, robust and reproducible profiling of the iPSC secretome is essential for dissecting extracellular signaling networks in human stem cell models and for advancing applications in disease modeling, drug discovery, and regenerative medicine.

Cell culture–based secretome analysis is a powerful approach for characterizing secreted proteins and cell-to-cell communication, providing valuable insights into disease biology and biomarker discovery. Mass spectrometry (MS)-based secretome proteomics has substantially expanded the ability to characterize extracellular proteins in cell culture systems^8–17^. Despite these advances, robust and reproducible secretome analysis in stem cells remains technically challenging. Secreted proteins are typically present at low abundance and can be readily obscured by residual media proteins or intracellular proteins released as a result of cell stress or damage^14,18^. Furthermore, key experimental parameters, such as supplement concentration, collection duration, cell density, sample clarification, and data-analysis workflows are often determined empirically and vary considerably across laboratories. For example, Hodgson-Garms *et al*.^12^ maintained mesenchymal stem cells in basal medium supplemented with insulin–transferrin–selenium for 48 h for secretome analysis, whereas other studies transitioned stem cells to serum-free or supplement-free media for 24 h to minimize contamination from exogenous media proteins^17,19^. However, conventional serum-starvation strategies may not be appropriate for stem cell systems, as stem cells require continuous growth factor supplementation to maintain pluripotency and viability. Therefore, the lack of a standardized workflow continues to limit reproducibility, cross-study comparability, and biological interpretation of stem cell secretome datasets.

A robust stem cell secretome workflow provides an opportunity to investigate how intracellular perturbations reshape extracellular signaling. Progranulin (PGRN), encoded by the *GRN* gene, is a secreted multifunctional glycoprotein that regulates cell survival, inflammation, and wound healing, while also playing a key role in lysosomal homeostasis^20–22^. Loss-of-function mutations in *GRN* cause frontotemporal dementia or lysosomal storage disease, depending on gene dosage^23^. We and others have previously shown that PGRN deficiency impairs lysosomal acidification, degradation, and trafficking across animal and human cell models^24–29^. Our recent multi-omics study further revealed intracellular molecular remodeling in both PGRN-deficient iPSCs and neurons and found that PGRN deficiency caused more perturbations of subcellular proteome in neurons compared to iPSCs^24^. However, how progranulin deficiency alters the extracellular proteome of iPSCs remains poorly understood. Investigating the secretome of isogenic *GRN*-knockout (KO) and wild-type (WT) iPSCs provides a unique opportunity to determine how progranulin deficiency remodels extracellular protein profiles.

In this study, we established an integrated iPSC secretome proteomics platform through systematic optimization of culture conditions, sample preparation, LC–MS acquisition, and downstream data analysis (**Figure 1**). The optimized workflow improves secretome coverage, quantitative reproducibility, and minimizes intracellular contamination while preserving stem cell health. We then applied this platform to isogenic *GRN*-KO and WT iPSCs to characterize disease-associated remodeling of the extracellular proteome, revealing coordinated changes in lysosome-associated secreted proteins. Together, this work provides a standardized framework for stem cell secretome proteomics and demonstrates its utility for studying extracellular signaling and disease-associated secretome remodeling in human stem cell models.

**Figure 1.**
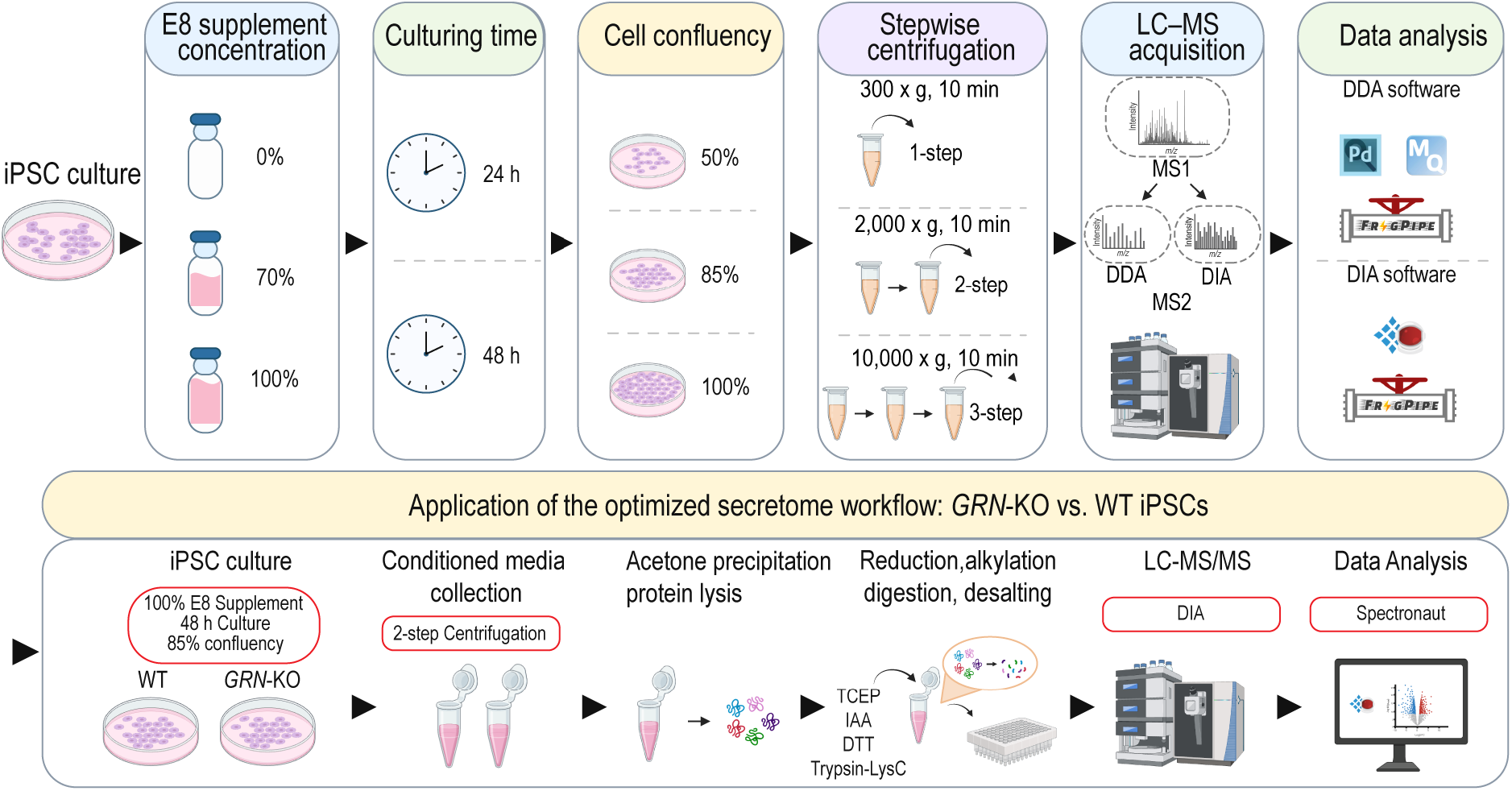
Schematic workflow of the optimized stem cell secretome proteomics platform. Optimized conditions used in the application experiment are highlighted in red. Portions of the figure were created in BioRender. Hao, L. (2026) https://BioRender.com/it2z2q9.

## 2. Experimental

### Human iPSC Culture and Media Harvest

Human iPSCs with stably expressed Dox-inducible NGN2 transcription factor were routinely cultured in our lab and maintained in Essential 8 (E8) medium (Gibco) in 10 cm dishes coated with Matrigel (Corning)^30^. For iPSC model of progranulin deficiency, isogenic WT and *GRN-*KO iPSCs were maintained in 6-well dishes (n=5). Accutase (Gibco) was used to dissociate iPSCs, and 10 μM ROCK inhibitor (ROCKi) was added to the media to prevent apoptosis until iPSC colony formation (typically for the first 24 h). Cells were then washed with phosphate-buffered saline (PBS) and maintained in complete E8 media without ROCKi. To optimize cell culture condition, iPSCs were washed three times with PBS to remove dead cells and incubated in E8 media with various concentrations of E8 protein supplement for 24 h or 48 h before collecting the media. The collected media were clarified by centrifugation to remove cell debris using three centrifugation protocols (**Figure 1**). The 1-step method involved a single centrifugation at 300 × g for 10 minutes. The 2-step method included an additional spin at 2,000 × g for 10 minutes. The 3-step method further added a third centrifugation at 10,000 × g for 10 minutes. After each centrifugation step, both cell pellet and supernatant were carefully separated and transferred in separate tubes for subsequent proteomics sample preparation.

### Proteomics Sample Preparation

All cell culture media samples and pellet samples were subjected to bottom-up proteomics sample preparation. Cell culture media samples from optimization experiments were processed by direct digestion in the collected and clarified media to avoid bias. Cell pellets from media collection centrifugation steps were resuspended in lysis buffer containing 8 M urea, 150 mM NaCl, and 50 mM ammonium bicarbonate. Protein lysate was sonicated for 15 min, and protein concentrations were determined using a detergent-compatible colorimetric protein assay (DCA, BioRad). Proteins from the *GRN*-KO vs. WT iPSC media were subjected to cold acetone precipitation, resuspension in urea lysis buffer, DCA assay, and normalization to the same total protein input. All protein samples were reduced with 5 mM tris(2-carboxyethyl)phosphine (TCEP) for 40 min, alkylated with 15 mM iodoacetamide (IAA) in the dark for 30 min, and quenched by 5 mM dithiothreitol (DTT) for 15 min to remove excess IAA. Trypsin/Lys-C was used for protein digestion at a 1:30 enzyme-to-protein ratio for 14 h, followed by an additional digestion at 1:60 ratio for 2 h at 37 °C with shaking at 1,200 rpm on a ThermoMixer. Digestion was quenched with trifluoroacetic acid (TFA) to pH<2. Peptides were desalted using a Waters Oasis HLB µElution plate and dried down by SpeedVac.

### LC-MS/MS Analysis

Two LC-MS systems were used in this study. Most samples from optimization experiments were analyzed on a Dionex UltiMate 3000 RSLCnano system coupled with a Thermo Scientific Q Exactive HF-X MS. *GRN*-KO vs. WT experiment was analyzed on a Neo Vanquish nanoUPLC coupled with a Thermo Scientific Astral MS. The mobile phase consisted of buffer A (2% acetonitrile, 0.1% formic acid in water) and Buffer B (0.1% formic acid in acetonitrile). For the Dionex LC platform, peptide samples were loaded onto an Acclaim PepMAP C18 trap column (3 μm, 100 Å, 75 μm × 2 cm) and subsequently separated on an Easy-spray PepMap C18 column (2 μm, 100 Å, 75 μm × 50 cm). The gradient was 210 min. Column temperature was maintained at 55 °C. LC flow rate was 0.25 µL/min. For the Neo LC platform, easy-spray PepMap C18 column (2 μm, 100 Å, 75 μm × 15 cm) without trap column was used. The gradient was 20 min. LC flow rate was 0.4 µL/min.

Both data-dependent acquisition (DDA) and data-independent acquisition (DIA) were conducted for secretome samples. For DDA, MS1 scans ranged from *m/z* 380 to 1,500 at 120,000 resolving power for Top-40 DDA. An automatic gain control (AGC) target was set at 1 × 10⁶ with a maximum injection time (maxIT) of 60 ms. The MS/MS analysis featured a resolving power of 7,500, an AGC target of 2 × 10⁵, and a 40 ms maxIT. Precursors were isolated within an *m/z* window of 1.4 and subjected to fragmentation using a normalized collision energy (NCE) of 30% and a dynamic exclusion time of 30.0 s. DIA analysis was performed using a precursor mass range of *m/z* 400–1000. MS1 spectra were acquired at 60,000 resolving power. MS1 settings included an AGC target of 1 × 10⁶ and a 60 ms maxIT. DIA MS/MS scans were acquired at 15,000 resolving power using an *m/z* 8.0 isolation window, a 27% NCE, a 1 × 10⁶ AGC target, and a 20-ms maxIT. Only DIA analyses were performed on Astral, using a precursor mass range of *m/z* 380–980. MS1 spectra were acquired at 240,000 resolving power. MS1 settings included an AGC target of 1 × 10⁶ and a 5 ms maxIT. DIA MS/MS scans were using Astral as the detector with an *m/z* 2.0 isolation window, 27% NCE, 3 × 10^4^ AGC target, and 5 ms maxIT.

### Proteomics Data Analysis

DDA raw data were analyzed using Proteome Discoverer (v 2.4.1.15, Thermo Scientific), MaxQuant (v 2.7.5.0)^31^, and FragPipe (v 24.0)^32^ software packages. DIA data were analyzed using the DirectDIA workflow in Spectronaut (v19.4, Biognosys) and the DIA-NN in FragPipe (v 24.0)^33^. The Swiss-Prot *Homo sapiens* reviewed database (2024_05 release) and our custom stem cell contaminant library were used for protein identification^34^. Search settings included Trypsin digestion allowing up to two missed cleavages, fixed carbamidomethylation of cysteine, and variable modifications for methionine oxidation and N-terminal acetylation. Peptide-spectrum matches (PSMs) and peptides were filtered by a false discovery rate (FDR) below 1%. Contaminant proteins were excluded from downstream analysis. DIA precursors with intensities below 1000 from the Q-Exactive HFX MS data were also removed.

### Statistical Analysis

Two-group statistical comparisons were performed using Welch’s t-test on log2-transformed values, with multiple-testing correction by the Benjamini–Hochberg (BH) procedure. Statistical significance in volcano plots was defined as adjusted-p value < 0.05 and |fold change| > 1.5. Data were presented as mean ± standard deviation (SD) from at least three biological replicates. All statistical analyses were conducted using GraphPad Prism 10 and R. Gene Ontology (GO) enrichment analysis was performed using ShinyGO (version 0.85.1)^35^.

### Fluorescence Microscopy Imaging

Human iPSCs were imaged regularly throughout the experiments to monitor morphology, cell density and signs of stress or detachment. For culture duration experiment, iPSCs stably expressing integrated green fluorescent proteins were imaged using the GFP channel, which specifically highlights the cytosol. Fluorescence imaging was performed using a CELENA® S digital imaging system (Logos Biosystems). All acquired images were processed and analyzed with ImageJ^36^.

## 3. Results and Discussion

### Full E8 Supplement Maximizes iPSC Health and Secretome Quality

Serum starvation is a common method for cell culture secretome. But many stem cell cultures are serum-free and require continuous growth factor supplementation to maintain pluripotency. Therefore, we first evaluated different concentrations of protein supplement in iPSC culture. Extensive cell death and detachment occurred with lower than 70% E8 protein supplement **(Figure 2A, Figure S1A)**. E8 supplement is well defined and contains only four proteins: insulin (INS), transferrin (TF), FGF2, and TGF-β1**(Figure 2B)**, allowing these components to be readily tracked in proteomic data. Compared with 70% supplement, 100% E8 yielded slightly more annotated secreted proteins and total protein identifications, more peptides per protein, and significantly less intracellular protein contamination (**Figure 2C, 2D, Supplementary Figure S1B–D**). Annotated secreted proteins were also more abundant with 100% E8 supplement (**Figure 2E, Supplementary Figure S1D**). Although the four E8 proteins were among the most abundant proteins detected, they did not appear to interfere with detection of other secreted proteins (**Figure 2F**). In contrast, supplement-free secretomes were dominated by intracellular contamination, with ACTB and GAPDH ranking as the two most abundant proteins. Thus, despite yielding more protein identifications, supplement-free condition primarily reflected intracellular leakage. These results support 100% E8 supplementation as the optimal condition for maintaining iPSC health and enabling robust secretome proteomic analysis.

**Figure 2.**
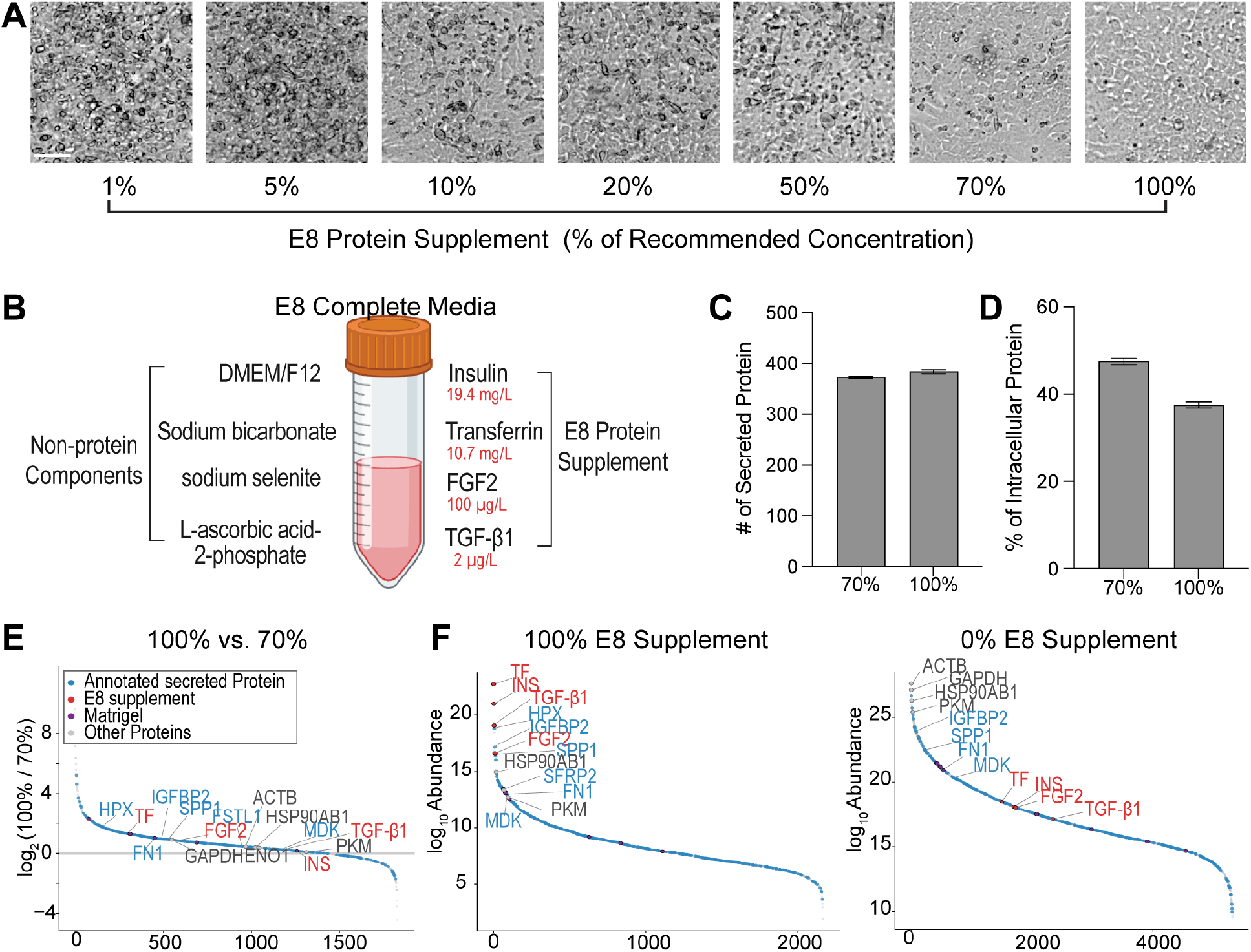
Effect of E8 protein supplement concentration on iPSC viability and secretome. **(A)** Bright-field microscopy images of iPSCs cultured with different percentages of E8 protein supplement. Scale bar denotes 50 μm. **(B)** Schematic summary of the complete E8 medium components. **(C)** Number of annotated secreted proteins quantified in conditioned media collected from iPSCs cultured with 70% and 100% E8 supplement. **(D)** Percentage of annotated intracellular proteins quantified from iPSC media with 70% and 100% E8 supplement. **(E)** Ranked protein abundance fold change from iPSC media between 100% and 70% E8 supplement conditions. **(F)** Ranked protein abundances from iPSC media with 100% E8 supplement (left) and 0% E8 supplement (right) conditions.

To identify and remove contaminants derived from E8 supplement, Matrigel coating, and common contaminants like trypsin and keratins, we created a stem cell-specific contaminant library (**Supplementary Table S1**, **Supplementary FASTA file**). This resource is freely available at https://github.com/HaoGroup-ProtContLib for incorporation into stem cell proteomics data analysis workflow.

### Optimization of iPSC Culture Duration and Confluency for Secretome Profiling

To further optimize the iPSC culture conditions for secretome analysis, we evaluated culture duration and cell confluency for collecting the conditioned media. Secretome profiles collected at 24 and 48 h after media change formed distinct principal component analysis (PCA) clusters (**Figure 3A, 3B**). The 48-h condition showed lower proteomic variability and greater proteome coverage (**Figure 3C, 3D**). Most annotated secreted proteins were detected at both time points, with 12 uniquely identified at 48 h (**Figure 3E**). Notably, volcano plot analysis revealed lower levels of several stress-associated proteins, including SERPINE1, HP, HPX, and APOH, at 48 h, whereas metabolically active proteins, including NAMPT, GPI, and the antioxidant enzyme SOD1, were higher at 48 h (**Figure 3F, 3G**). These findings suggest that iPSCs require time to recover from media-change-associated stress and support a 48-h collection period for reducing proteomic variability and improving secretome coverage.

**Figure 3.**
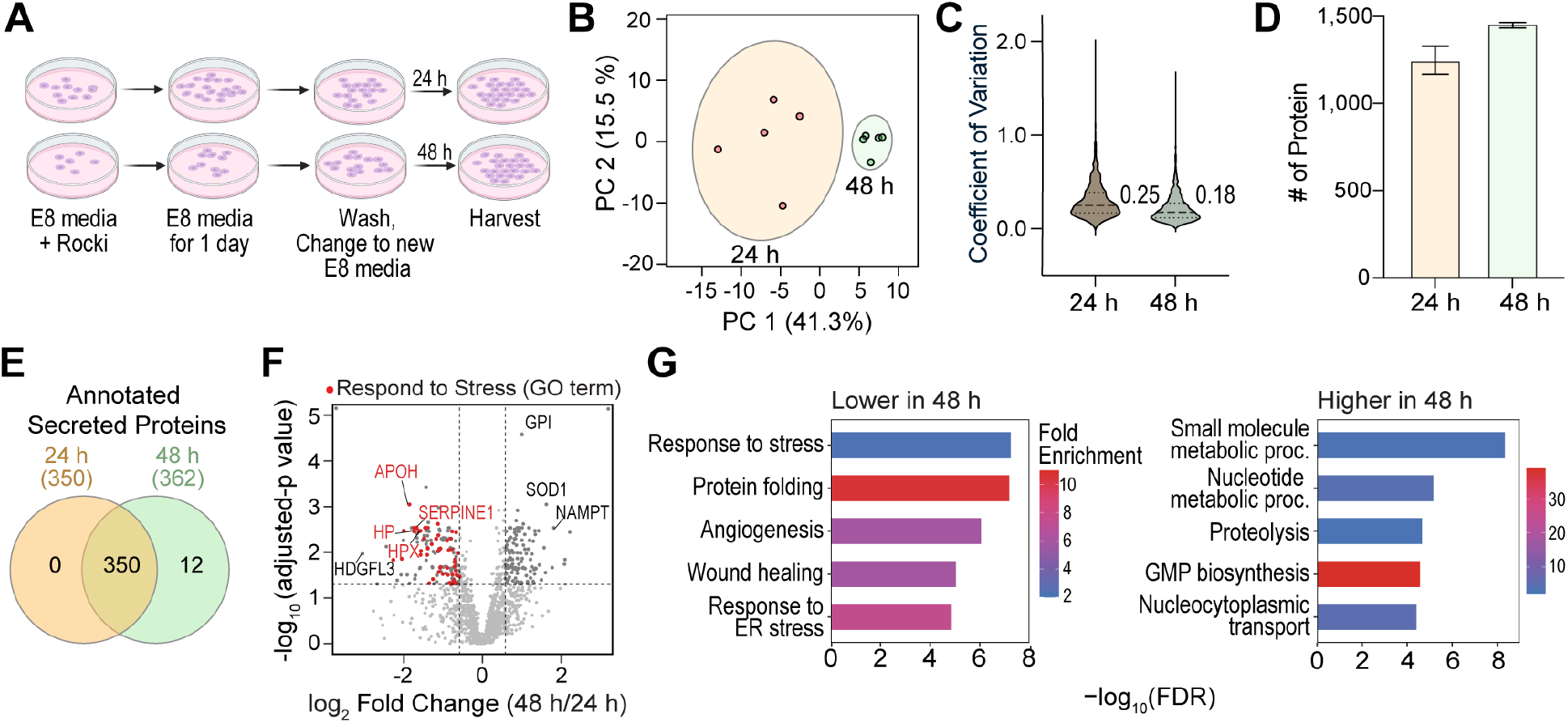
Effect of culture duration on iPSC secretome proteomics. **(A)** Experimental workflow comparing iPSC media collected after 24 h or 48 h of fresh media change. **(B)** PCA plot of secretome proteomics profiles from 24 h or 48 h of fresh media change. **(C)** Distribution of protein CVs across biological replicates; median CV values are indicated on the graph. **(D)** Total number of proteins quantified after 24 h and 48 h of conditioned media collection (N = 5; mean ± SD). **(E)** Venn diagram showing the overlap of annotated secreted proteins quantified from the two groups. **(F)** Volcano plot comparing protein abundances between the 48 h and 24 h secretomes. Proteins annotated with the GO term response to stress are highlighted in red. **(G)** GO enrichment analysis showing biological processes from proteins significantly lower (left) and higher (right) in 48 h group compared to 24 h group.

To assess the impact of cell density on secretome profiling, we compared iPSCs at 50%, 80%, and 100% confluency at the time of media collection by seeding them at different densities (**Supplementary Figure S2A**). Overall, secretome profiles were largely comparable across conditions, with similar numbers of quantified proteins and peptides and slightly greater proteome coverage at 80% and 100% confluency (**Supplementary Figure S2B-G**). PCA and volcano plot analyses further showed highly similar proteomic profiles between the 80% and 100% conditions. To minimize cellular stress and overgrowth-associated spontaneous differentiation, 80% confluency was selected as the optimal condition for secretome collection.

### Optimization of Media Clarification by Stepwise Centrifugation

To remove cell debris, conditioned media are typically clarified by centrifugation prior to secretome analysis. However, centrifugation protocols vary considerably across studies in speed, duration, and number of steps, with no standardized approach for secretome sample preparation. We therefore compared three stepwise centrifugation strategies and analyzed both the clarified media and resulting pellets by proteomics (**Figure 4A**). PCA showed that secretomes from 1-step centrifugation clustered separately from the 2- and 3-step conditions, whereas the latter showed similar profiles (**Figure 4B**). One-step centrifugation yielded the most protein identifications (**Figure 4C, 4D**), but uniquely detected proteins were enriched for mitochondrial and organelle membrane components, indicating greater cellular contamination (**Figure 4E**). These results indicate that a single centrifugation at 300 g for 10 min, widely used in secretome studies, can remove most cell debris but still contain substantial cellular proteins in the clarified media.

**Figure 4.**
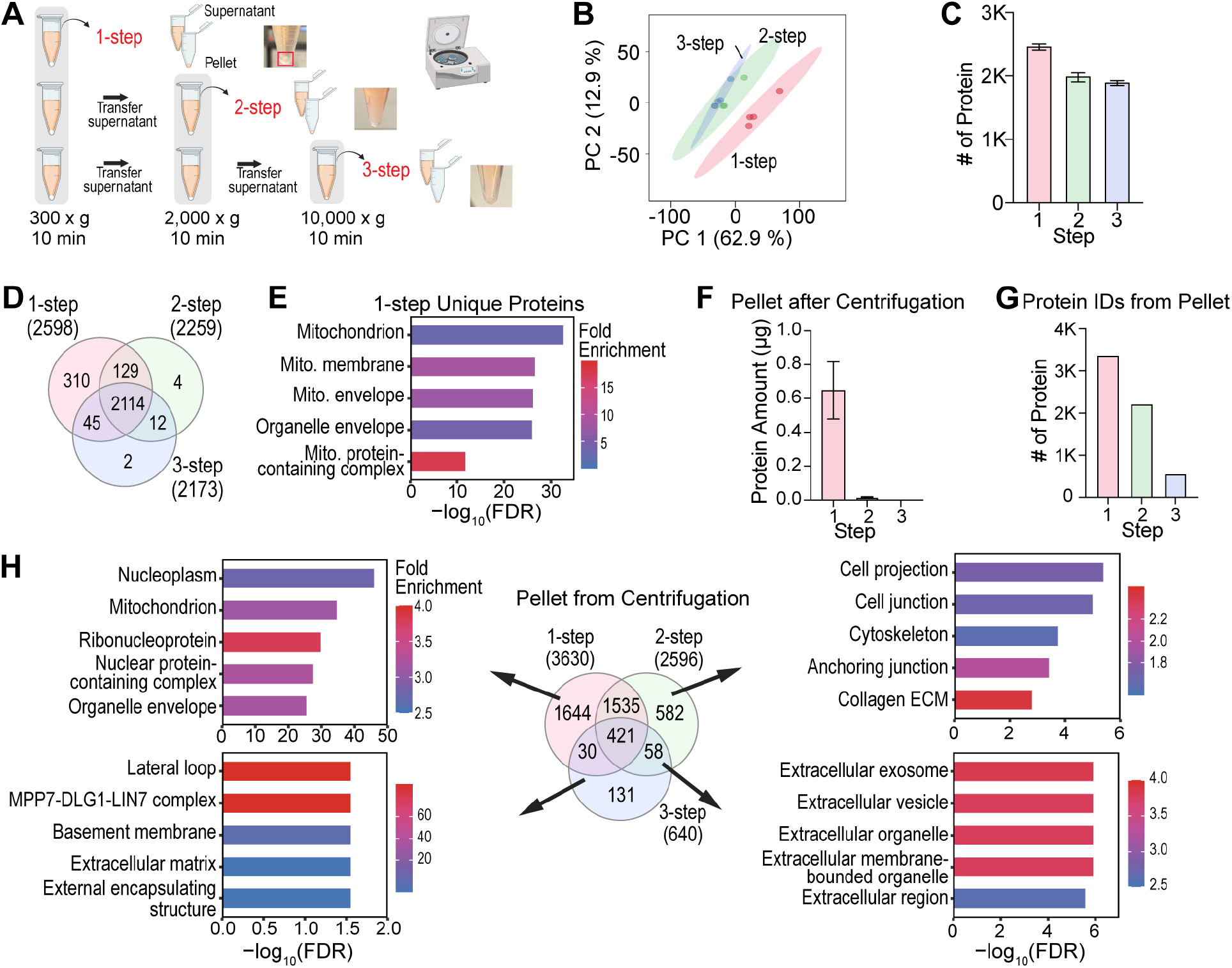
Effect of centrifugation strategies on secretome protein recovery and intracellular contamination. **(A)** Schematic workflow of stepwise centrifugation for iPSC media clarification. **(B)** PCA plot of secretome profiles obtained using three centrifugation methods. **(C)** Total number of proteins identified for each method. (**D)** Venn diagram showing total protein overlap across the three methods. **(E)** Enriched cellular component GO-terms from 310 unique proteins identified in 1-step centrifugation. **(F)** Total protein amount recovered in the cell pellet after each centrifugation step, measured by colorimetric protein assay. **(G)** Number of proteins identified from pellets collected after each centrifugation workflow. **(H)** Venn diagram comparing proteins identified from pellets recovered from the three centrifugation methods as well as the enriched cellular component GO-terms.

Analysis of the centrifugation pellets further supported this conclusion. As expected, the first-step pellet contained substantially more protein and protein identifications than subsequent pellets (**Figure 4F, 4G**) and was enriched for intracellular components (**Figure 4H**). The second-step pellet was enriched for cell projection, cell junction, cytoskeletal, and extracellular matrix (ECM) proteins, suggesting further removal of residual cellular material but minor loss of ECM proteins. In contrast, the third-step pellet was enriched for extracellular vesicle- and exosome-associated proteins, suggesting potential loss of biologically relevant extracellular components. Although the second centrifugation step also removed some extracellular proteins, it substantially reduced cellular contamination while preserving greater secretome coverage compared to other methods. Therefore, 2-step centrifugation was selected as the optimal media clarification strategy for iPSC secretome analysis.

### Evaluation of LC-MS/MS Acquisition and Proteomics Data Analysis Strategies

For LC-MS/MS-based secretome proteomics analysis, we compared DDA and DIA LC-MS/MS acquisition methods and analyzed the raw data with several commonly used data analysis platforms, including MaxQuant, FragPipe, and Proteome Discoverer (PD) for DDA data and FragPipe with DIA-NN and Spectronaut for DIA data. PD and Spectronaut provided the highest number of total protein and annotated secreted protein identifications for DDA and DIA data, respectively (**Figure 5A, 5B**). FragPipe DDA showed the lowest variation of protein abundances across all DDA and DIA platforms (**Figure 5C**). Overall, DIA outperformed DDA regarding proteome coverage and peptide-per-protein counts (**Figure 5A, D, E**). But the coverage of annotated secreted proteins was comparable across workflows, indicating that both DDA and DIA platforms provide robust proteomics quality for secretome analysis (**Figure 5E**). Thus, while all workflows provided robust secretome profiling, DIA-Spectronaut was selected for downstream biological analyses due to its comprehensive coverage and reproducibility. Together, these experiments established the optimized conditions for iPSC secretome profiling, with the impact of key optimization parameters summarized in **Supplementary Figure S3**.

**Figure 5.**
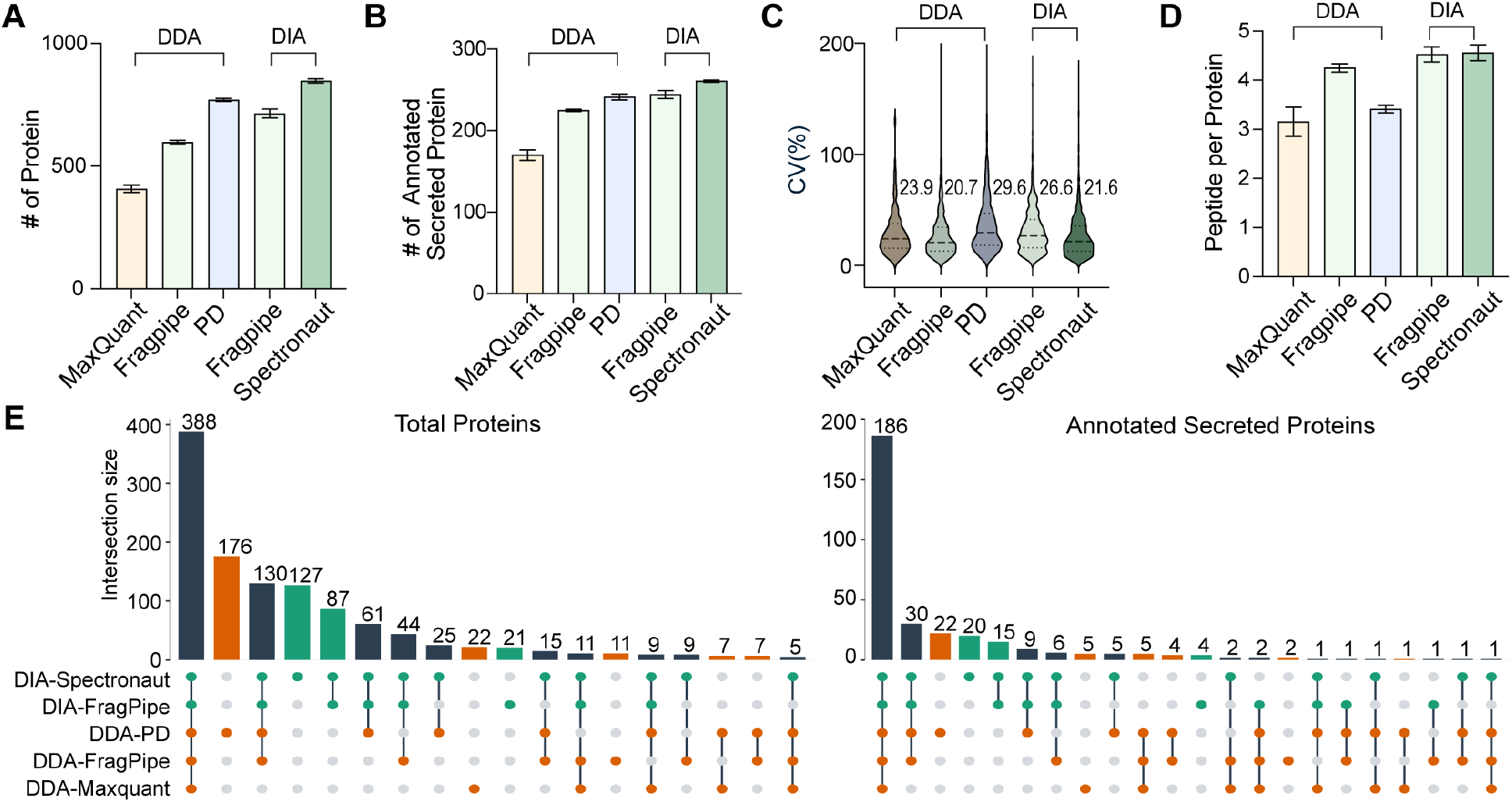
Comparison of DDA and DIA LC-MS/MS method and data analysis platforms for secretome proteomics. **(A)** Total protein identifications obtained from DDA data analyzed with MaxQuant, FragPipe, and Proteome Discoverer, and from DIA data analyzed with FragPipe and Spectronaut. **(B)** Number of annotated secreted proteins identified using each workflow. **(C)** Distribution of protein CVs across workflows. Median CV values are indicated. **(D)** Average number of peptides identified per protein for each analysis workflow. (**E**) UpSet plot showing the overlap of protein identifications across five workflows for total proteins (left) and annotated secreted proteins (right).

### Progranulin Deficiency Alters the iPSC Secretome

With the optimized secretome workflow established, we next applied it to an iPSC disease model to investigate how progranulin deficiency reshapes the extracellular proteome in comparison with the intracellular proteome (**Figure 6A**). PCA showed complete separation of secretomes from isogenic *GRN*-KO and WT iPSCs (**Figure 6B**). Progranulin was readily detected in WT iPSC media but was absent in *GRN*-KO media samples (**Figure 6C**). Differential expression analysis revealed widespread remodeling of the secretome caused by progranulin deficiency (**Figure 6D**). Notably, multiple lysosomal hydrolases and lysosome-related proteins, such as CTSC, CTSZ, CTSL, LIPA, MANBA, PPT1 were significantly down-regulated in *GRN*-KO iPSC secretome. GO enrichment analysis of downregulated proteins revealed enriched biological processes related to wound healing, response to growth factor, angiogenesis and negative regulation of protein metabolic processes **(Figure 6E)**. These downregulated pathways agreed with the known extracellular functions of progranulin in growth factor–like signaling, tissue remodeling, and cellular stress responses^37^. The coordinated reduction of extracellular lysosomal hydrolases represents a previously unrecognized consequence of progranulin deficiency and prompted comparison with intracellular proteomic alterations.

**Figure 6.**
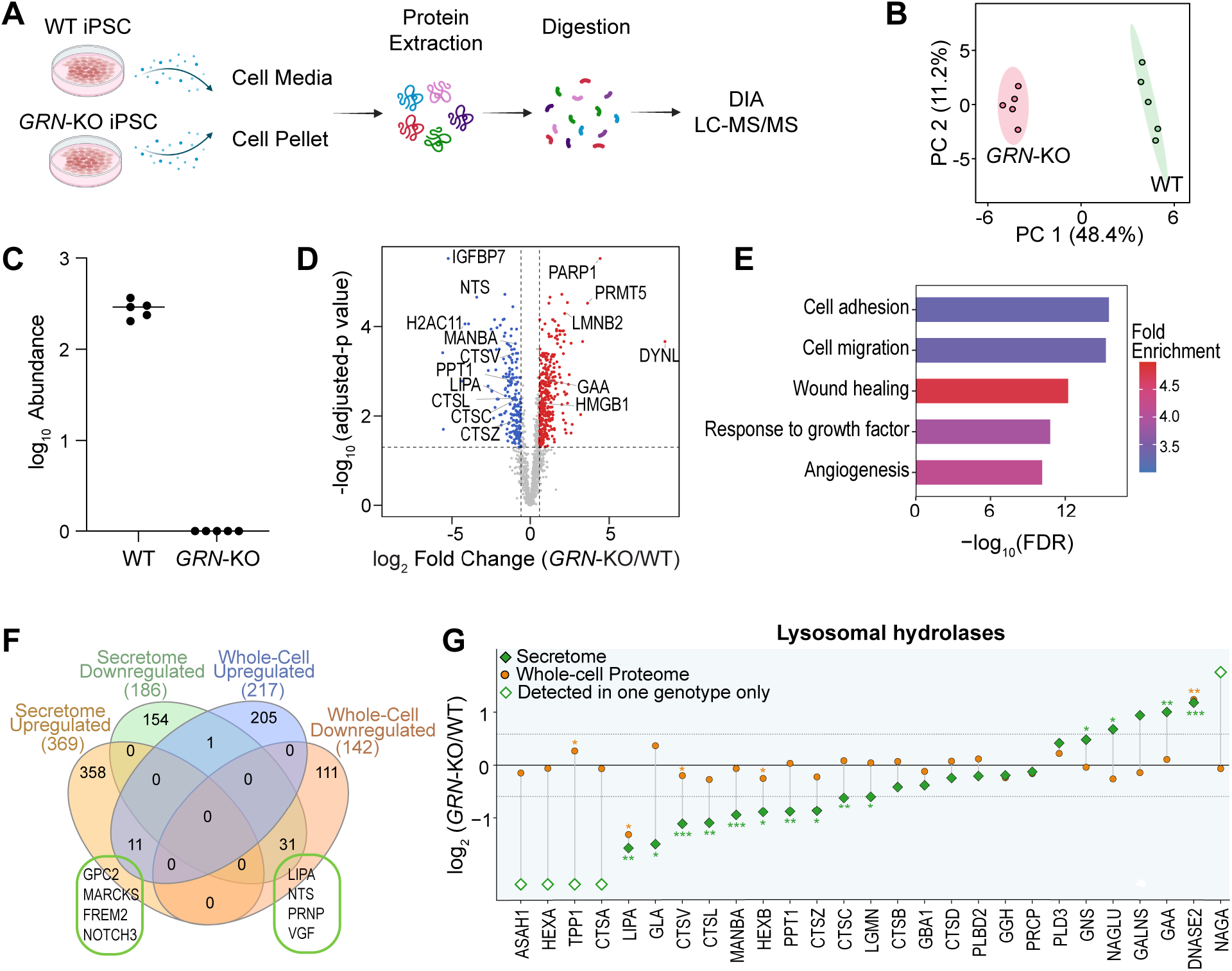
Secretome proteomics from progranulin-deficient iPSC model. **(A)** Schematic of the experimental workflow comparing secretomes from WT and *GRN*-KO iPSCs (N=5). **(B)** PCA plot of WT and *GRN*-KO secretome profiles. **(C)** Progranulin protein abundance quantified in WT and *GRN*-KO iPSC secretomes. **(D)** Volcano plot comparing *GRN*-KO vs. WT secretome proteomics. **(E)** Enriched biological process GO-terms from proteins significantly decreased in the *GRN*-KO vs. WT comparison. **(F)** Venn diagram comparing proteins significantly altered in the *GRN*-KO vs. WT iPSC secretome and cellular proteomics. **(G)** Relative abundance changes of key proteins in the secretome and cellular proteomics of *GRN*-KO vs. WT iPSCs.

To determine whether alterations in the secretome reflect or diverge from intracellular proteome changes, we compared the significantly altered proteins identified in the GRN-KO secretome with those changed in the *GRN*-KO whole-cell lysate proteome from our previous study^24^ (**Figure 6F**). Overall, progranulin deficiency produced distinct proteomic alterations in the extracellular and intracellular compartments. Whereas numerous lysosomal hydrolases were consistently reduced in the *GRN*-KO secretome, their intracellular abundance showed more heterogeneous changes. Notably, TPP1, CTSD, and PLD3 exhibited opposite trends, with reduced extracellular abundance but increased intracellular abundance in *GRN*-KO cells^24^ **(Figure 6G)**.

We and others have previously demonstrated that progranulin deficiency causes accumulation of lysosomal hydrolases within neuronal lysosomes together with impaired lysosomal degradative capacity and elevated lysosome pH in neurons^24–29^. Lysosomal hydrolases are constitutively secreted through incomplete mannose-6-phosphate sorting and subsequently retrieved by the secretion–recapture pathway^38–40^, making the extracellular abundance of these proteins a useful indicator of lysosomal trafficking. Because progranulin itself is delivered to lysosomes through sortilin and partially M6P-dependent routes^41,42^, loss of progranulin may broadly disrupt lysosomal protein trafficking. Therefore, the combination of reduced extracellular lysosomal hydrolases and their intracellular accumulation may suggest impaired lysosomal trafficking as well as a possible reduction in lysosomal exocytosis in *GRN*-KO iPSCs.

## 4. Conclusions

In this study, we established a standardized workflow for iPSC secretome proteomics through systematic optimization of culture conditions, sample preparation, LC–MS acquisition, and data analysis. The optimized workflow which used full-strength E8 supplemented medium, a 48 h media collection period, 80% cell confluency, two-step centrifugation, and DIA-LC–MS/MS improved secretome coverage, quantitative reproducibility, and minimized intracellular contamination while preserving stem cell health. Application of this workflow to isogenic *GRN*-knockout iPSCs revealed extensive extracellular proteome remodeling, including a coordinated reduction of lysosomal hydrolases that was not reflected in the intracellular proteome. Together with previous evidence of lysosomal hydrolase accumulation and lysosomal dysfunction in progranulin-deficient cells, these findings suggest altered lysosomal protein trafficking and a possible reduction in lysosomal exocytosis. Although distinguishing bona fide secreted proteins from proteins released through extracellular vesicles, unconventional secretion, or cellular leakage remains challenging, this optimized workflow provides a robust framework for stem cell secretome proteomics and enables mechanistic investigation of extracellular proteome remodeling in disease models.

## Supplementary Information

- Supplementary Figure S1: Optimization of E8 supplement percentages for iPSC secretome.
- Supplementary Figure S2: Effect of cell plating density on iPSC secretome.
- Supplementary Figure S3: Summary of the effects of key optimization parameters on stem cell secretome quality.
- Supplementary Table S1: Stem cell-specific contaminant protein list.
- Supplementary Table S2: Secretome proteomics results of GRN-KO vs. WT iPSCs.
- Supplementary file: Stem cell-specific contaminant FASTA library.

## Author Contributions

J.N. and L.H. designed the study. J.N. conducted iPSC cell culture and fluorescence imaging. J.N. and H.T. conducted proteomic experiments and data analysis. J.N. conducted LC-MS analysis. J.N. wrote the first draft of the manuscript with revisions from L.H. and H.T. All authors have read and agreed to the final version of the manuscript.

The authors declare no competing financial interest.

## Data availability

All MS raw files have been deposited to MassIVE with the project accession ID MSV000101126.

## Acknowledgements

This study is supported by the NIH grants (R01NS121608). The authors would like to thank Noah Smeriglio and Dr. Gwangbin Lee from the Hao Lab for maintaining the LC-MS instruments and all other Hao Lab members for their feedback and support.

## TOC

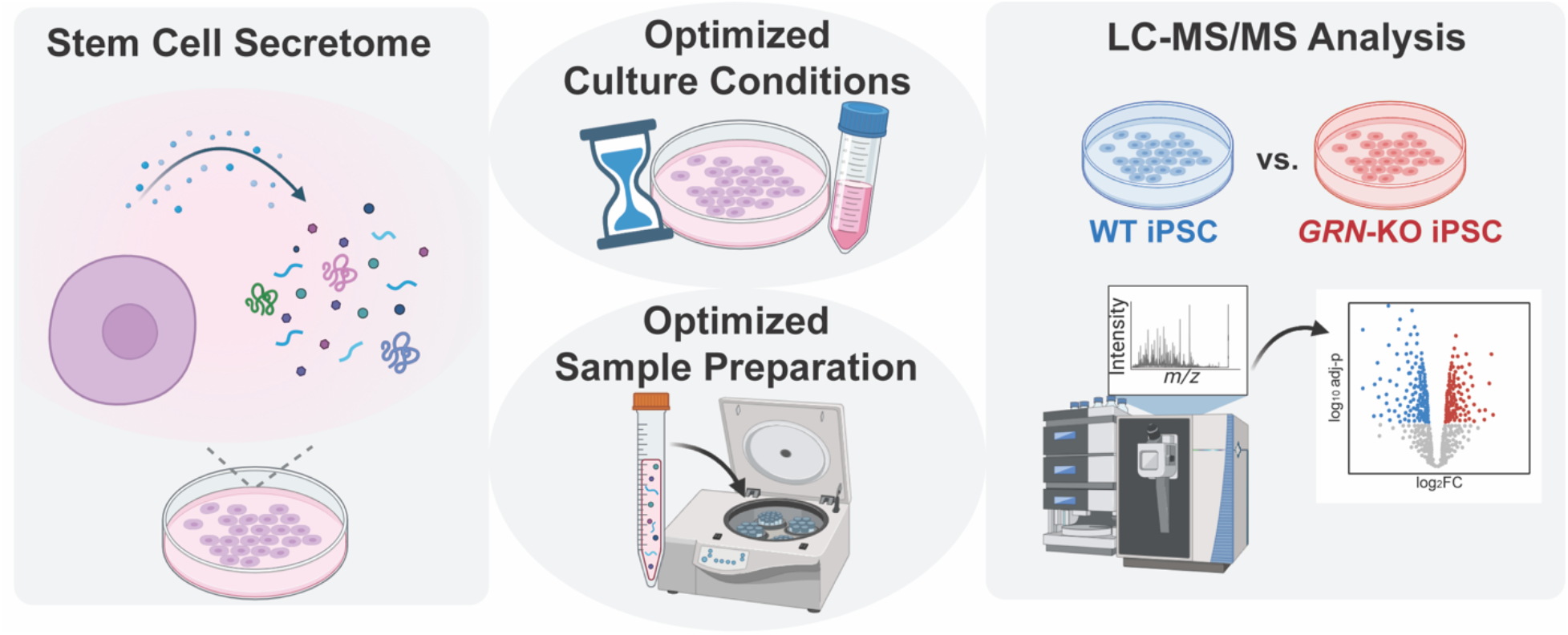

## Notes

### Competing Interest Statement

The authors have declared no competing interest.

