## Supplemental Information for "An Optimized Stem Cell Secretome Proteomics Platform: Application to Progranulin-Deficient iPSCs"

*Corresponding author

Ling Hao, PhD.

Associate Professor

Department of Chemistry & Biochemistry

University of Maryland

### Table of Contents


- Supplementary Figure S1: Optimization of E8 supplement percentages for iPSC secretome.
- Supplementary Figure S2: Effect of cell plating density on iPSC secretome.
- Supplementary Figure S3: Summary of the effects of key optimization parameters on stem cell secretome quality.
- Supplementary Table S1: Stem cell-specific contaminant protein list.
- Supplementary Table S2: Secretome proteomics results of *GRN*-KO vs. WT iPSCs.
- Supplementary file: Stem cell-specific contaminant FASTA library.

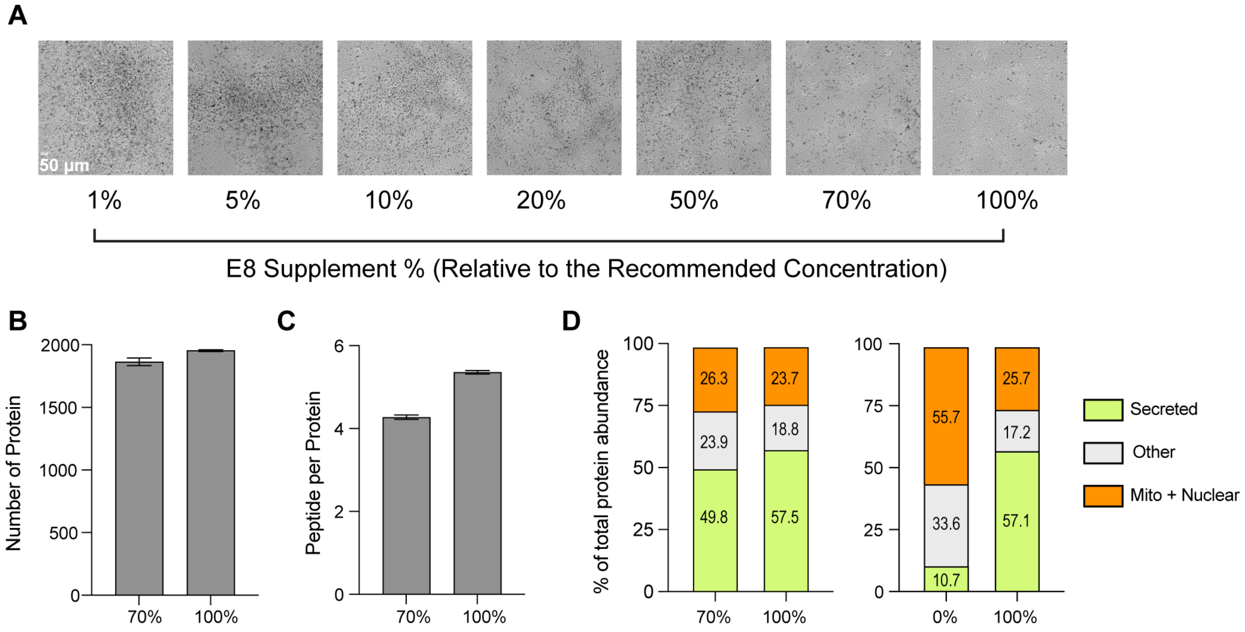


**Supplementary Figure S1. Optimization of E8 supplement percentages for iPSC secretome. (A)** Additional representative bright-field microscopy images of iPSCs cultured with increasing E8 protein supplement concentrations Scale bar denotes 50 µm. **(B)** Total quantified proteins from iPSCs media with 70% and 100% E8 supplement. **(C)** Average number of peptides quantified per protein with 70% and 100% E8 supplement. **(D)** Relative percentage of proteins classified as annotated secreted, mitochondrial and nuclear proteins as representative cellular proteins, or other proteins under different E8 supplement concentrations.

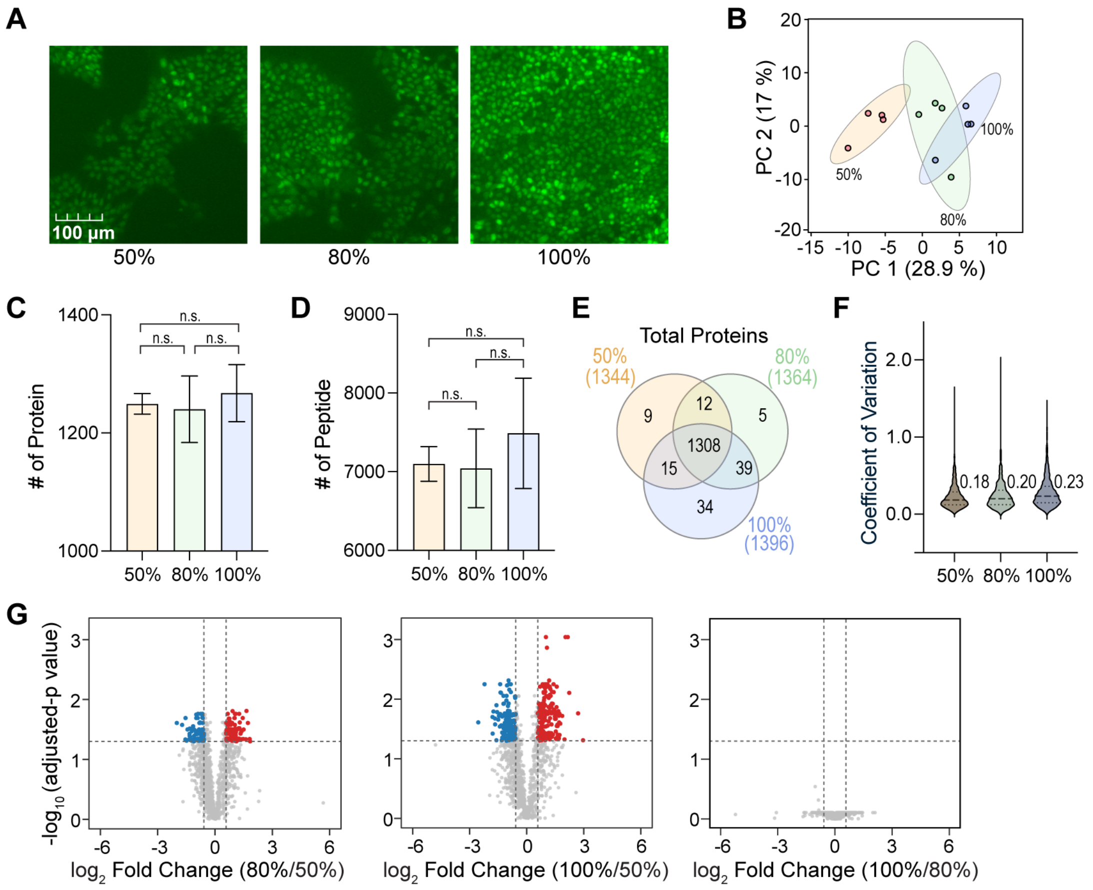


**Supplementary Figure S2. Effect of cell plating density on iPSC secretome. (A)** Fluorescence microscopy images of iPSCs seeded at 50%, 80%, and 100% confluency. Cells stably expressing green fluorescent protein (GFP) were used for visualization. Scale bar denotes 100 µm. **(B)** PCA plot of secretome profiles from the three plating densities. **(C)** Total protein identifications from iPSC secretome at each plating density. **(D)** Total peptide identifications from iPSC secretome at each plating density. **(E)** Venn diagrams showing overlap of total proteins among the three plating densities. **(F)** Distribution of protein CVs across biological replicates for each plating density. **(G)** Volcano plots of secreted proteomics from 80% vs. 50% and 100% vs. 50% confluency, and 100% vs. 80% comparisons.

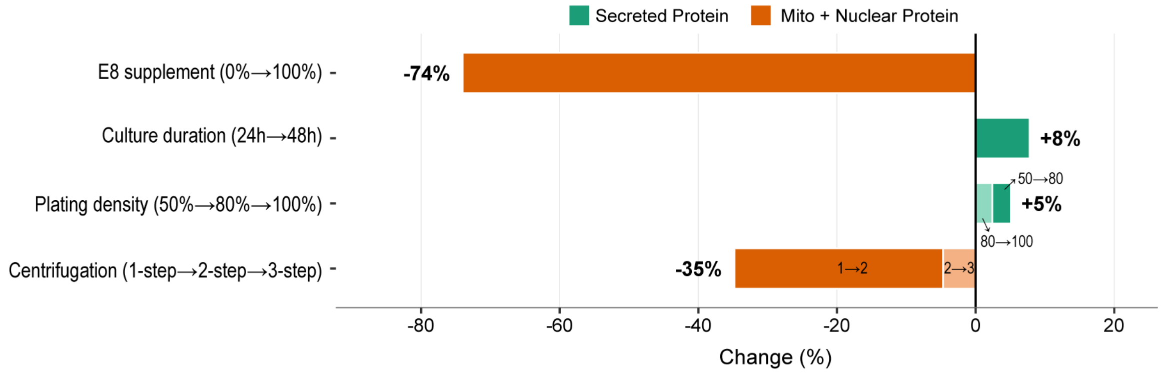


**Supplementary Figure S3. Summary of the effects of key optimization parameters on stem cell secretome quality.** Labels within the bars indicate the corresponding change between conditions. For example, changing E8 supplementation from 0% to 100% reduced the relative abundance of mitochondrial and nuclear proteins by 74% (representing example cellular proteins). Changing culturing duration from 24 h to 48 h increased the relative abundance of annotated secreted proteins by 8%.
